# Auto-SMART: a web-based platform for repair-template design for CRISPR/Cas9 homology-directed editing

**DOI:** 10.64898/2026.09.16.752223

**Authors:** Chuanping Zhao, Yan Cao, Kirill A. Martemyanov

**Author notes:** Corresponding authors: Dr. Kirill Martemyanov Dr. Chuanping Zhao.

## Abstract

Genome engineering via CRISPR/Cas9 has been widely applied in biological research and developing gene therapies. However, genome editing flexibility is strongly constrained by the position and quality of guide RNAs relative to the intended edit: desired edit sites are frequently positioned far from efficient protospacer adjacent motifs (PAM). SMART (Silently Mutate And Repair Template) technique relaxes this constraint by silently mutating the gap between the Cas9 cut and the edit in the repair donor, preserving the encoded protein while expanding the usable guide-RNA design space. Designing such donors manually requires coordinated reasoning over transcript structure, reading frame, candidate protospacers, codon usage, homology arms and the expected edited allele. Auto-SMART is a web platform that automates this process for CRISPR/Cas9-mediated insertion, deletion, point mutation and sequence replacement in coding exons and in introns. Users choose a genome assembly, search a gene symbol or transcript, select a modification site and select an editing operation. Auto-SMART finds nearby *Streptococcus pyogenes* Cas9 (SpCas9) guides, ranks them by predicted on-target activity and cut-to-edit geometry, builds repair templates with synonymous substitutions, automatically recodes residual on-target sites to prevent re-cutting and renders interpretable panels for the target locus, donor template and locus after HDR, together with a downloadable report and a Primer3 genotyping handoff. The web application for Auto-SMART is freely available at https://auto-smart.chuanping.org/.

## Introduction

CRISPR/Cas9 has made targeted modification of endogenous loci routine ^1,2^, yet the efficiency of homology-directed repair remains strongly dependent on the position of guide RNA relative to the intended modification site ^3,4^. For knock-ins, deletions and precise substitutions, the desired editing site is often not adjacent to an efficient 5’-NGG-3’ PAM. As the distance between the double-strand break and the modification site grows, the repair template increasingly fails to drive the desired change ^3,5^. This limits the efficiency of point mutagenesis, epitope insertion and other precise modifications.

SMART editing addresses this constraint at the level of the repair template design. In a SMART donor, the sequence between the cut site and the desired edit, named gap sequence, is silently mutated so that it no longer base-pairs perfectly with the cut chromosome, while the encoded amino acid sequence is preserved ^5^. This biases repair toward incorporation of the full template and makes it possible to position guides farther from the edit site, thereby broadening the practical design space. SMART has been validated for efficient *in vivo* labeling of endogenous proteins in the vertebrate retina and the brain ^5^. However, manual design of a SMART templates by hand is cumbersome and error-prone: it requires mapping the edit onto the coding frame, enumerating candidate protospacers, choosing synonymous codons consistent with the organism’s codon usage, assembling homology arms with correct geometry, checking that the repaired allele will not simply be re-cut, and finally designing genotyping primers.

Existing web applications cover individual elements of this workflow. CRISPOR provides comprehensive guide discovery, specificity and activity scoring ^6,7^; genome browsers supply transcript and sequence context ^8,9^; codon-usage resources support synonymous codon choice ^10,11^; and Primer3 and its interfaces design PCR and sequencing primers ^12–14^. However, users need to switch between individual resources and even then, none currently provides SMART template designs.

Auto-SMART is a browser application built to close this gap. It performs SMART template design and returns output organized around the practical needs: guide selection, mapping cut and editing sites, template bases to be silently changed, and sequence to synthesize, expected allele after modification, how to block re-cut and genotype edited allele. Both coding exons and introns of the selected transcript can be targeted, with intronic designs confined to a splice-safe interior so that the donor and acceptor signals of the intron are not altered.

## Results

### Design principle and processing pipeline

Auto-SMART converts a selected modification site and an intended edit into a synthesis-ready SMART design. With a conventional template, knock-in efficiency decreases steeply once the site to be edited moves beyond ∼10 bp from the cut site. This largely occurs due to undesired base pairing between the gap sequence in the donor template and broken target locus, restoring the original sequence instead of installing the edit ^3,5^. SMART eliminates that competition: silently recoding the gap prevents it from pairing with the target, keeps the entire donor engaged, and maintains knock-in efficiency at approximately half the optimal level even when the gap is ∼100 bp ^5^. This design is automatically implemented in Auto-SMART at codon resolution. Every codon overlapping with the gap is replaced by the synonymous codon closest in usage frequency to the native one based on the species-specific codon usage table (Figure 1A).

**Figure 1.**
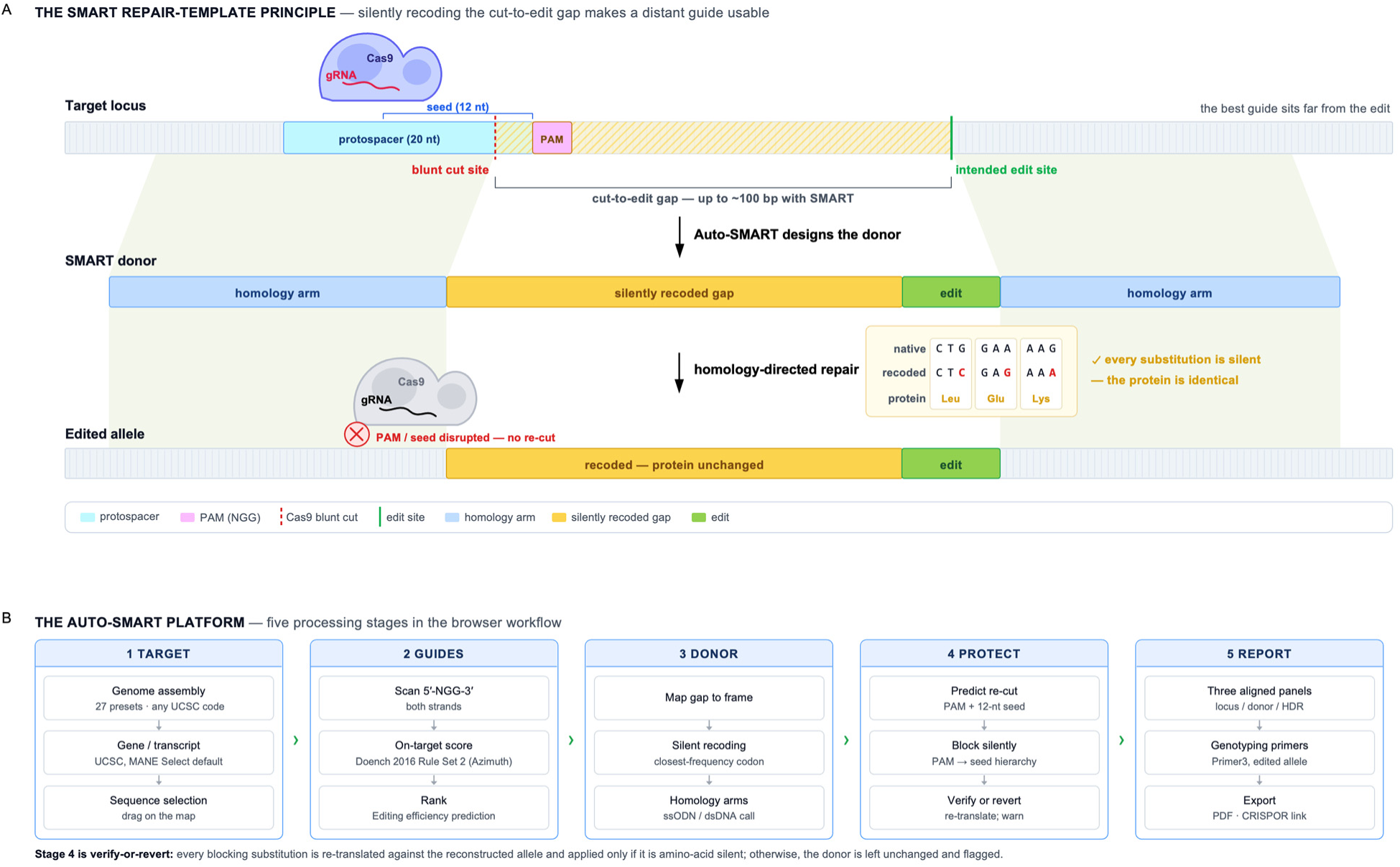
The SMART design principle and the Auto-SMART processing pipeline. (A) The genome editing principle that Auto-SMART automates. A Cas9 cut guided by a distant, high-activity protospacer lies up to ∼100 bp from the intended edit; in the donor, the cut-to-edit gap is silently recoded and flanked by homology arms, and the inset shows the reading frame preserved codon by codon (native and recoded codons encode the same amino acids). After homology-directed repair the edited allele carries the intended edit and resists re-cutting because the PAM or seed has been silently disrupted. (B) The five processing stages of the server: TARGET (assembly, transcript, Sequence selection), GUIDES (NGG scan on both strands, Doench Rule Set 2 scoring, ranking by predicted editing efficiency), DONOR (gap mapping, closest-frequency synonymous recoding, homology arms and ssODN/dsDNA call), PROTECT (re-cut prediction from PAM and 12-nt seed, silent blocking, verify or revert) and REPORT (three aligned panels, Primer3 primers seeded from the edited allele, PDF and CRISPOR links).

The application runs the design in five stages (Figure 1B). The output of each stage can be directly examined. The first stage, TARGET renders the gene as a curated transcript and fixes the selected sequence as the coordinate anchor for subsequent analysis. The second stage, GUIDES positions SpCas9 guides on both strands near the selected sequence and ranks them by predicted on-target activity discounted by cut-to-edit distance, so that a distant but active guide can outrank a close but weak one. The third component, DONOR assembles the designated gap, the edit and the homology arms into one template and determines whether it is suitable for synthesis as an ssODN or dsDNA donor. Fourth, PROTECT introduces synonymous mutations into the PAM or guide seed sequence to prevent edited alleles from being recleaved by SpCas9 complexes. Finally, REPORT returns the result card described below (Figure 2B), together with a downloadable report and CRISPOR link for each guide.

**Figure 2.**
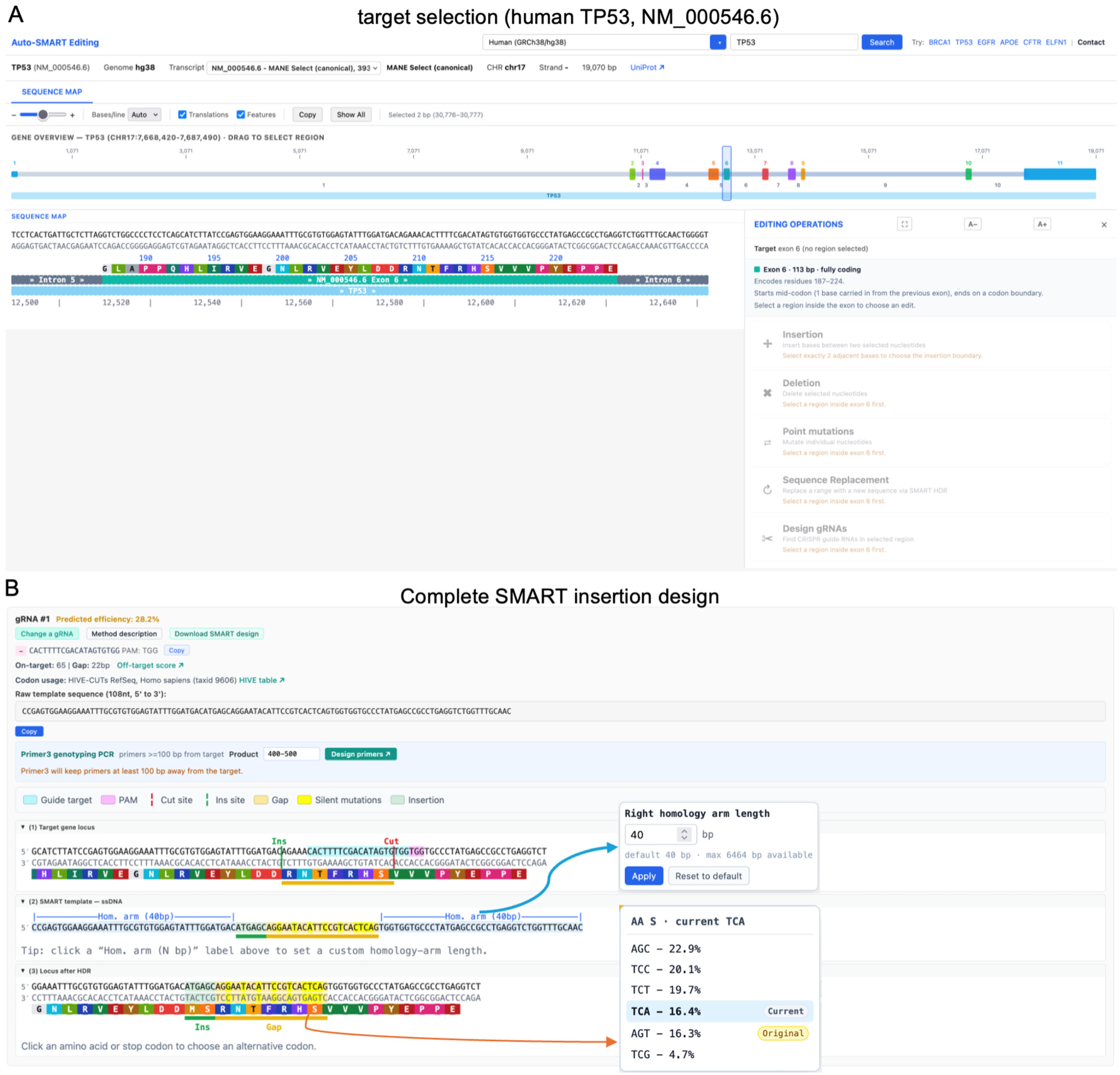
The Auto-SMART interface and a complete SMART design for a representative target. (A) Target selection in reference to coding sequences: the sequence map shows nucleotide, exon and translation tracks; dragging across a sequence of interest exposes the four editing applications and a guide-only option in the editing-operations panel. (B) A complete SMART insertion result card: the summary header reports the guide, PAM, predicted on-target score, cut-to-edit gap and codon-usage source; the raw donor sequence and a Primer3 genotyping handoff are provided directly; and three aligned panels show the target locus, the SMART donor template and the locus after HDR. Insets: homology-arm length can be set independently for each side (upper), and clicking any residue opens the synonymous codons available for it with their usage frequencies, marking both the current and the native codon (lower).

### Integrated workspace

Auto-SMART organizes the entire design into one browser workspace. After choosing an assembly and searching a gene or transcript, the user sees a sequence map with nucleotide, exon and translation tracks (Figure 2A). Dragging across a sequence exposes the editing operations whose requirements match the selection. For example, insertion needs an adjacent two-nucleotide boundary, whereas deletion, point mutation and sequence replacement act on selected nucleotides. Guide-only option is also available. The selected sequence remains the coordinate anchor for all downstream processes, obviating the need to copy coordinates or sequence when transitioning between different stages.

Rather than providing a fixed output, the result card is made auditable and editable (Figure 2B). Homology arm length can be set per side, and clicking any amino acid in the edited allele opens the full set of synonymous codons for that residue, each annotated with its usage frequency in the selected table, with the current and the native codon both marked. Every such change re-runs re-cut prediction and re-renders all three panels.

### Four editing applications originating from a single selection

We designed an application for the same workspace to support four most frequent CRISPR/Cas9 applications: insertion, deletion, point mutation and sequence replacement (Figure 3). Insertion places a user-supplied sequence at the chosen position (Figure 3A). When the insertion site occurs immediately upstream of the stop codon, Auto-SMART switches automatically to the C-terminal SMART-RC/CT geometry (Figure 3B). This geometry has both homology arms anchored at the cut site and the native stop codon is retained downstream. Deletion removes the selected nucleotides (Figure 3C). Point mutation changes one or more selected bases (Figure 3D). Finally, sequence replacement swaps the selected sequence for a new sequence of equal or different length (Figure 3E). In each case, Auto-SMART scans nearby guides, ranks them, recodes the gap where applicable, and returns a result card (Figure 2B). As an illustrative example, we used the human TP53 MANE Select transcript (NM_000546.6, GRCh38/hg38) in exon 6 to generate insertion, deletion, point-mutation and sequence-replacement designs (Figures 2 and 3), each providing the guide/PAM, cut position, gap length, predicted activity, raw donor and primer handoff.

**Figure 3.**
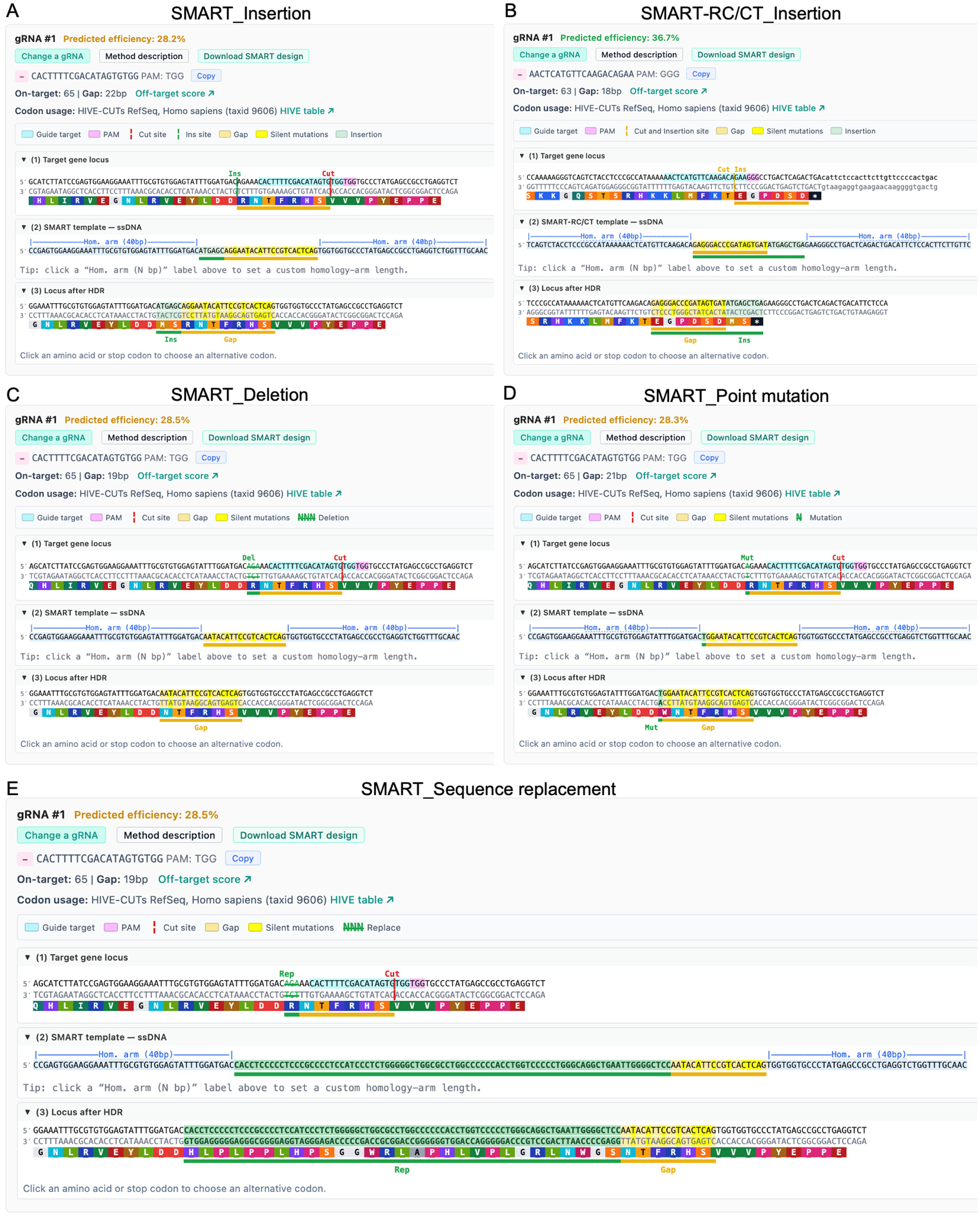
SMART designs for the various editing applications. Result cards for representative designs: (A) SMART insertion; (B) SMART-RC/CT insertion, used automatically when the insertion sits immediately upstream of the stop codon, in which both homology arms are anchored at the cut and the native stop is retained downstream; (C) SMART deletion; (D) SMART point mutation; (E) SMART sequence replacement. Each card reports a ranked guide and shows the relationship among guide, PAM, cut site, silent gap mutations, donor template and the expected locus after HDR, with the translated protein displayed beneath every panel.

### Automatic amino-acid-silent re-cut avoidance

One caveat of CRISPR/Cas9 editing occurs when the repaired allele retains a cut-competent site. In order to mitigate it, we designed Auto-SMART to detect when this occurs and neutralize them with synonymous edits, highlighting the change and confirming in the HDR panel that the protein is unchanged. It distinguishes three outcomes: PAM block (Figure 4A), seed block (Figure 4B) and unresolvable (Figure 4C). To quantify this behavior in the current build, we classified the top-ranked design in each of 1,152 coding windows across 36 human genes (Figure 4D). We found that 14.8% of designs retained a residual on-target site after HDR. Of these, Auto-SMART silently neutralized 99.4% (38 by disrupting the PAM and 132 by recoding the seed) and flagged the remaining single case as predicted re-cut rather than altering the protein. In no case was a non-synonymous change introduced to avoid re-cutting.

**Figure 4.**
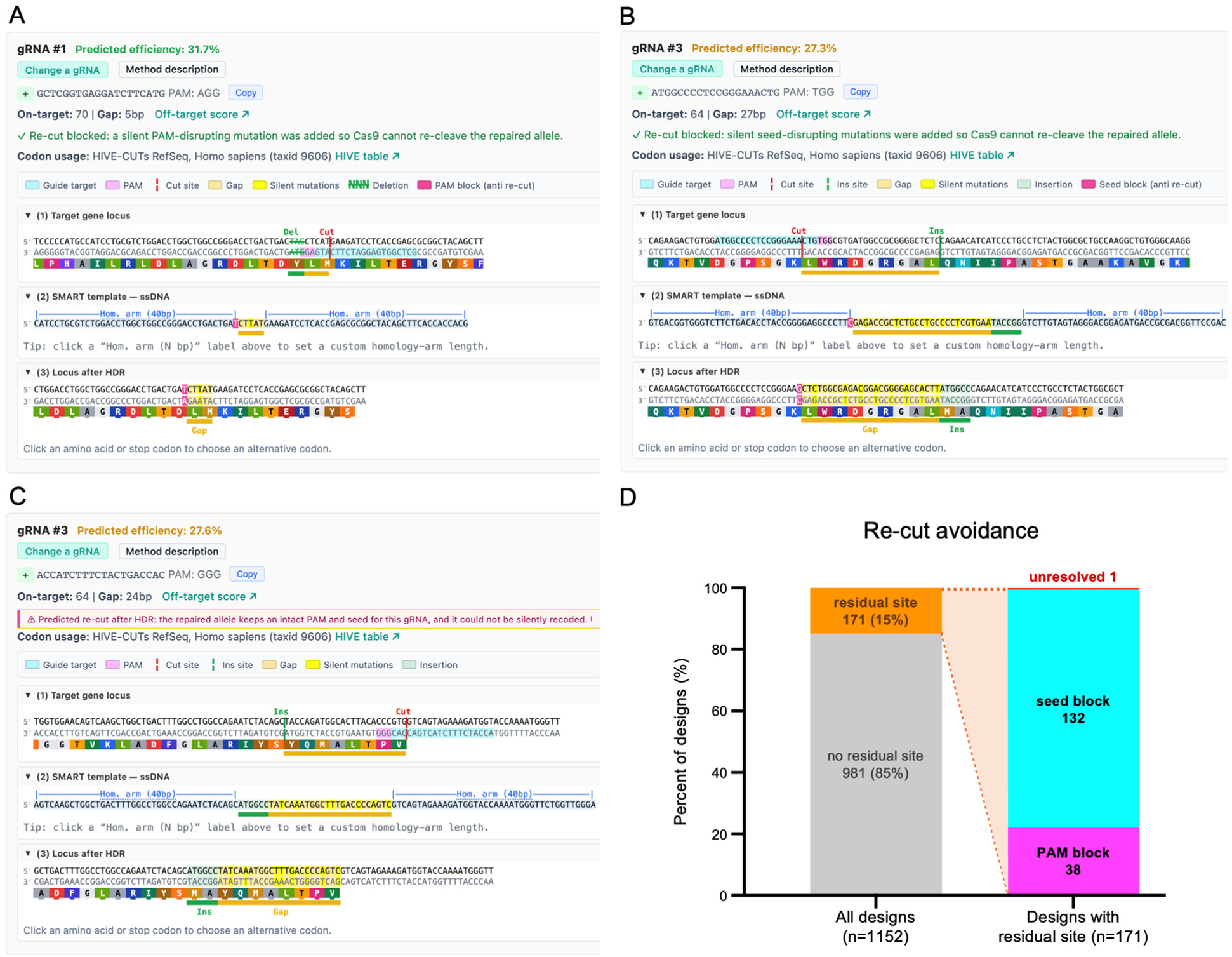
Automatic amino acid silent re-cut avoidance. The three outcomes Auto-SMART distinguishes when a repaired allele would remain cut-competent. (A) PAM block: a silent PAM-disrupting substitution (highlighted) prevents Cas9 from re-cleaving the edited allele. (B) Seed block: where the PAM cannot be broken silently, synonymous substitutions are placed in the PAM-proximal seed instead. (C) Unresolvable: no silent change removes the risk, so the design is returned unchanged with an explicit “predicted re-cut” warning rather than altering the protein. (D) Frequency of each outcome across 1,152 top-ranked designs in 36 human genes.

### Intron editing with splice-site protection

Short cargo such as recombinase sites is routinely placed in introns. In such cases, the insert is spliced out and need not preserve a reading frame. Auto-SMART allows every intron as a target (Figure 5A), reporting length and terminal dinucleotides and withholding the splice signals (25 bp donor, 60 bp acceptor), leaving an editable interior. Human TP53 intron 1 (10,754 bp, GT..AG) yields a 10,669 bp interior with 1,496 splice-safe SpCas9 guides (Figure 5A).

**Figure 5.**
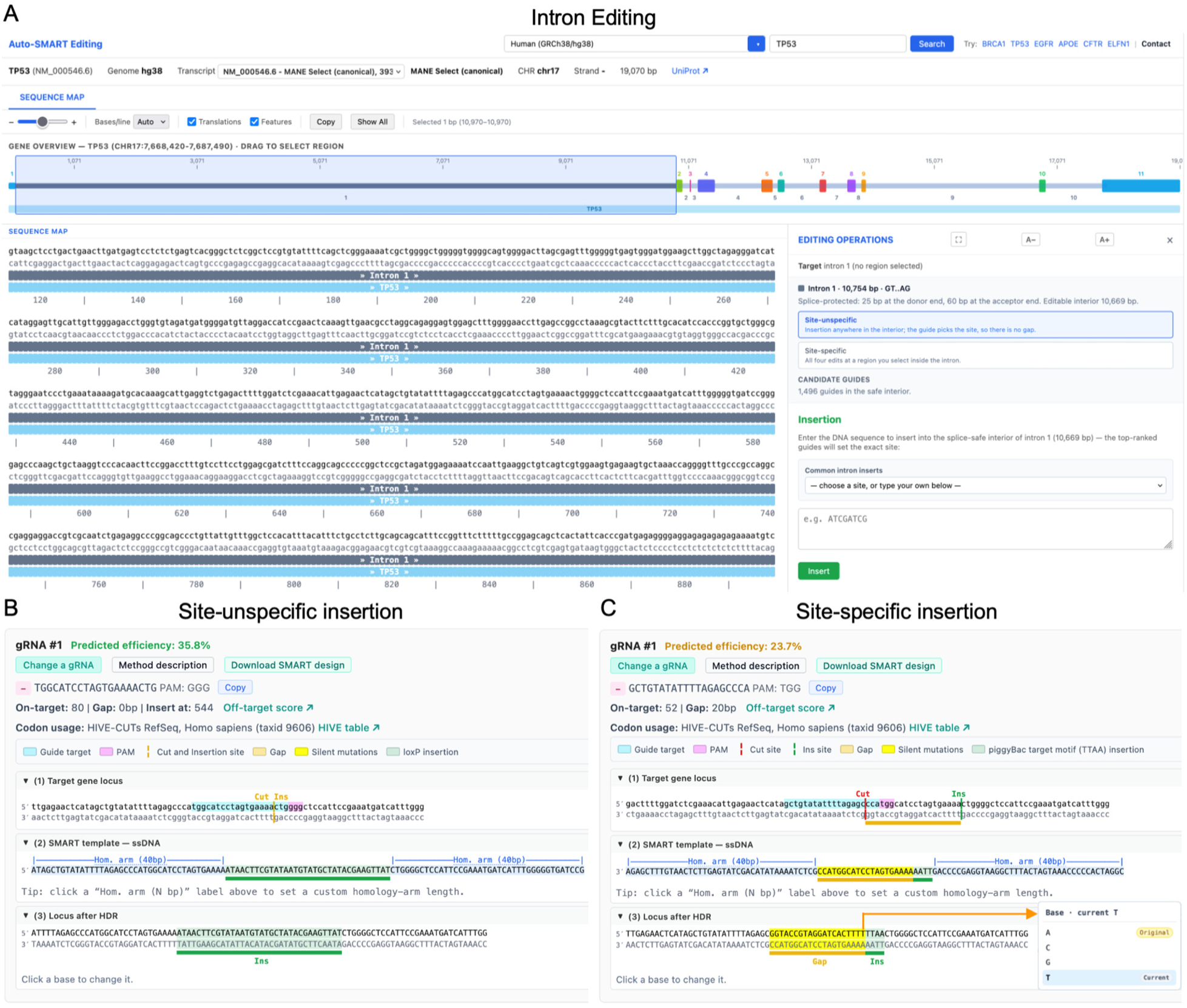
Intron editing with splice-site protection. (A) Selecting an intron activates a dedicated design mode (human *TP53*, NM_000546.6; hg38). For intron 1 (10,754 bp; GT–AG), 25 bp at the donor end and 60 bp at the acceptor end are protected to preserve splice-site function. The remaining 10,669-bp editable interior contains 1,496 splice-safe SpCas9 guides. In site-unspecific mode, the top-ranked guide determines the insertion site. In site-specific mode, the user selects a target region for insertion, deletion, point mutation, or sequence replacement. (B) Site-unspecific insertion of a 34 bp loxP site: the cut and insertion sites are same, so the gap is 0 bp and no recoding is needed. (C) Site-specific insertion of the piggyBac TTAA motif 20 bp from the cut; the gap is recoded base by base (yellow) to block re-annealing and re-cutting. Inset: with no protein encoded, the codon picker becomes a base picker.

We incorporated two options in the design. For the insertion at unspecified site, a guide’s cut site determines where the insertion site is, making the cut-to-edit gap zero, and restricting ranking to on-target activity. For example, a 34 bp loxP in intron 1 returns a 114 nt ssODN around the best guide (on-target 80, 35.8% predicted efficiency; Figure 5B). Alternative site-specific mode runs all four applications as in an exon, but the gap is recoded by complementation rather than synonymous substitution, changing every base, leaving no intact protospacer or PAM, and preserving GC content (Figure 5C).

### Coverage, throughput and transcript fidelity

We benchmarked Auto-SMART across a panel of 20 vertebrate and model-organism assemblies and the four editing applications (Figure 6A, B). All 2,400 design tasks (30 genes x 4 applications x 20 assemblies) completed successfully, including the 10 assemblies whose codon-usage tables were fetched live rather than bundled (Figure 6A). The design time was well under one second per task, with median completion time of 154-284 ms across the four applications (Figure 6B). Insertion was the slowest likely because the SMART-RC/CT geometry required extra checks for the longer template. Per-assembly medians were the shortest for S. cerevisiae (54 ms) and longest for human (354 ms). Across 600 genes per species, translations of Auto-SMART’s default transcript matched the reviewed UniProt canonical sequence for 91.0% of human genes and 83.0% of mouse genes, rising to 91.2% and 85.1% when restricted to RefSeq Select transcripts (Figure 6C). The discordances were predominantly isoform/length differences arising from RefSeq-versus-UniProt annotation choices rather than translation errors, and each was flagged for the user at design stage (Figure 6D).

**Figure 6.**
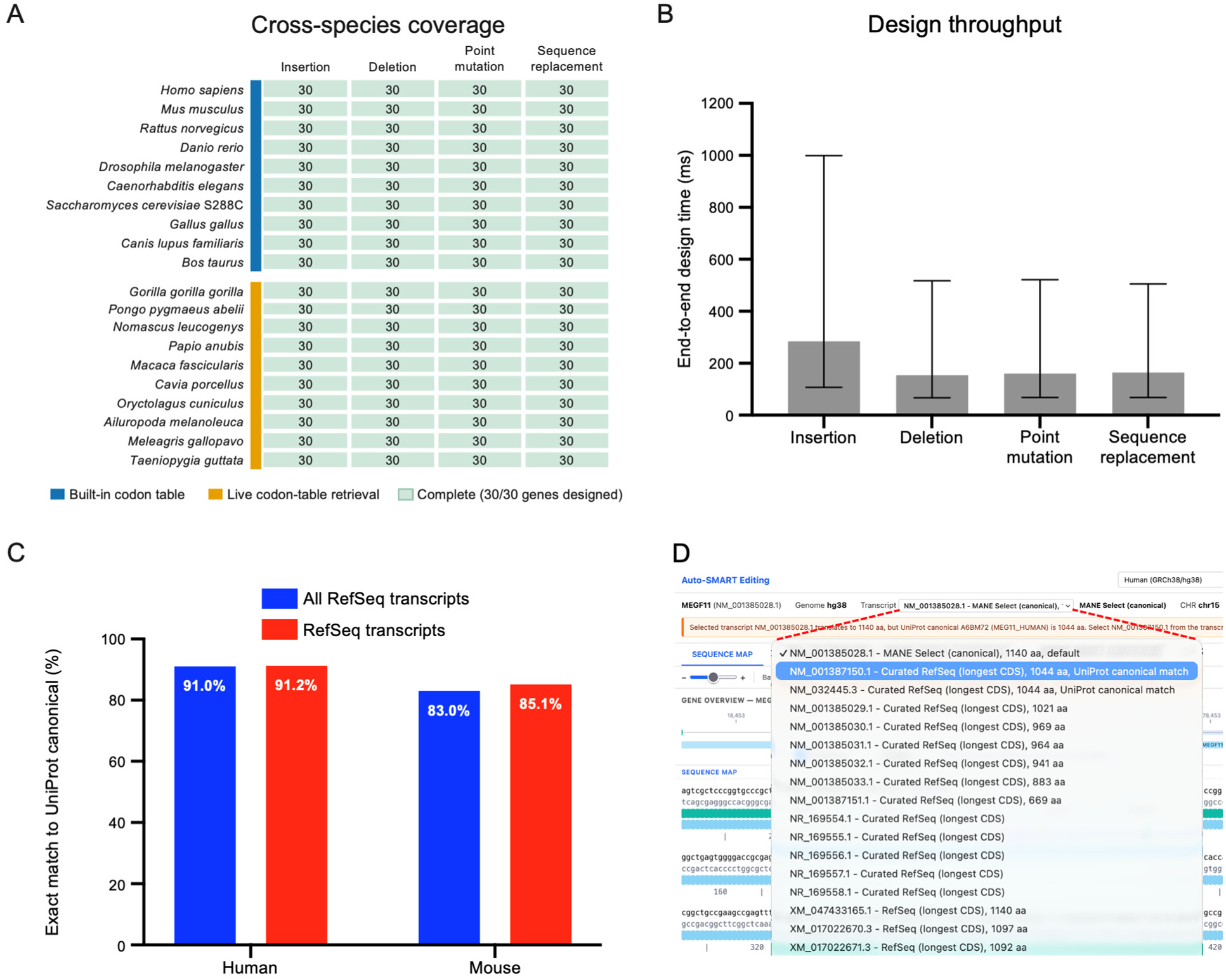
Validation of Auto-SMART. (A) Cross-species coverage: across a 20-assembly panel and four applications, all 2,400 design tasks completed (cell values are the number of distinct genes designed). (B) End-to-end design throughput (bars, median; whiskers, 25th-75th percentile; n = 600 designs per application). (C) Concordance between the protein translated from Auto-SMART’s default transcript and the reviewed UniProt canonical sequence for 600 comparable genes per species, for all RefSeq transcripts and for RefSeq Select only. (D) Where the two disagree, the transcript dropdown lets the user switch to the isoform matching the UniProt canonical protein, so the annotation choice is explicit at design time.

### SMART expands the usable guide space

The benchmarks above establish that Auto-SMART produces complete designs quickly and comprehensively. However, they do not address the magnitude effect enabled by the SMART technique: silently recoding the cut-to-edit gap allows using distant guides. Thus, we next measured the extent of the extra guide space provided by Auto-SMART sampling. Across 4,000 coding edit sites in 100 human genes and 37,980 scored SpCas9 guides, 91.4% of coding positions have at least one usable guide that satisfies the conventional constraint of keeping the edit within 10 bp of the cut. This number drops to only 44.2% if an additional criterion is introduced requiring such a guide to be predicted as highly active (Rule Set 2 score >= 60 ^15,16^; Figure 7A). Widening the window to the 100 bp that SMART supports raises those figures to 99.8% and 93.8%, while increasing the median number of usable guides per site from 3 to 19, a ∼6-fold gain (Figure 7B). This also increases the median best achievable on-target score from 59 to 69 (Figure 7C), suggesting that SMART does not only allow wider guide selection but also increases their quality. The effect is uniform across loci: guides were improved for all 100 examined genes, with a median gain of 47.5% per-gene from mean of 44.2 +/- 13.9% to 93.8 +/- 8.5%.

**Figure 7.**
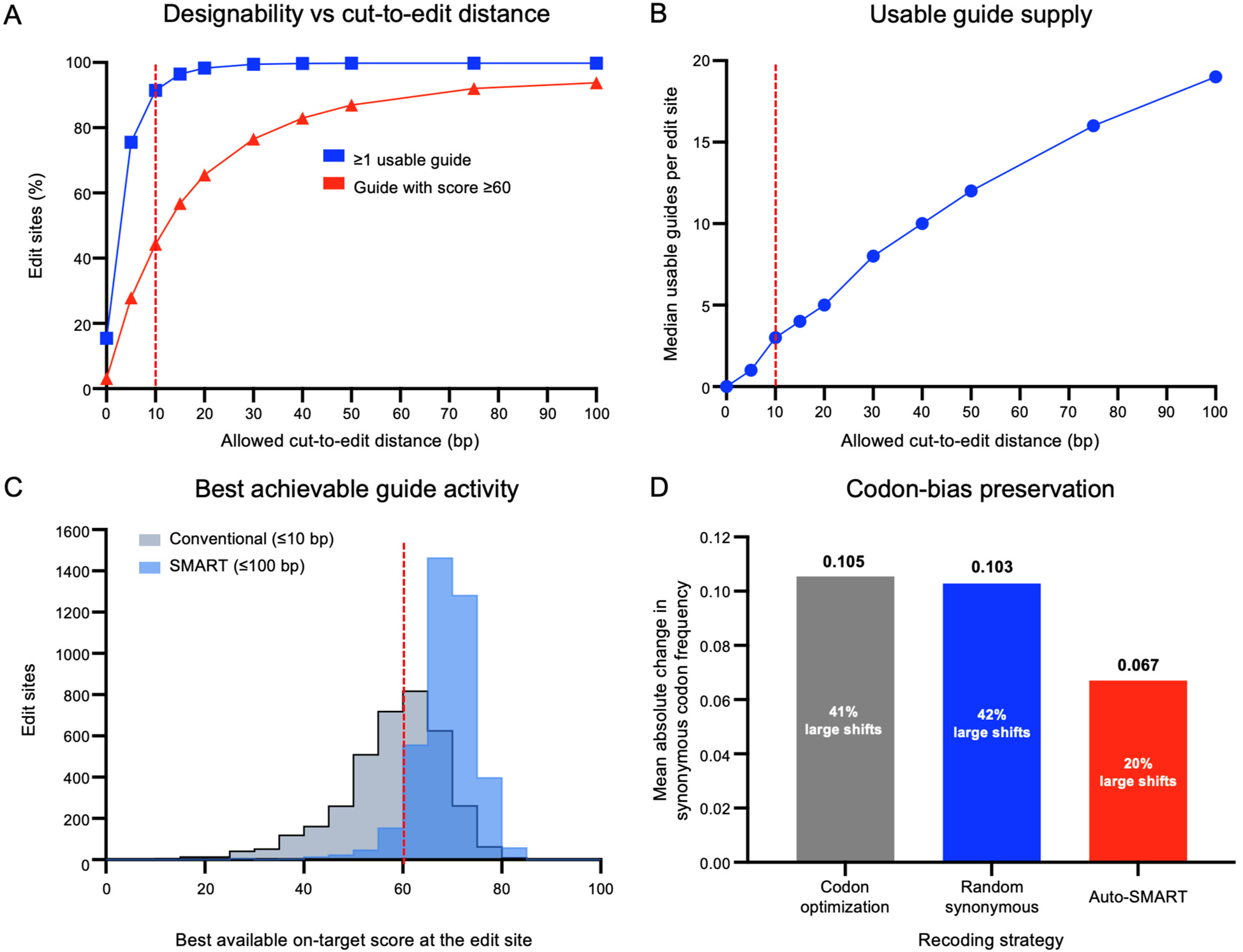
SMART expands the usable guide space. Measured over 4,000 coding edit sites in 100 human genes (40 evenly spaced sites per gene) and 37,980 scored SpCas9 guides. (A) Proportion of edit sites with at least one usable guide, and with a guide predicted to be highly active (Rule Set 2 >= 60), as a function of the allowed cut-to-edit distance; the conventional 10 bp constraint is marked. (B) Median number of usable guides per edit site against allowed distance. (C) Distribution of the best on-target score achievable at each edit site under the two windows. (D) Codon-bias preservation: mean absolute change in relative synonymous codon usage produced by codon optimization, by random synonymous choice and by Auto-SMART’s closest-frequency rule, over the 59 re-codable human codons.

Because the donor should perturb native codon usage as little as possible, we further compared the closest-frequency rule with codon optimization and with random synonymous choice across all 59 human codons that have a synonymous alternative (Figure 7D). The mean absolute change in relative synonymous codon frequency was 0.067 for closest-frequency recoding, 0.105 for optimization and 0.103 for random choice, a 36% reduction that persisted after weighting each codon by its abundance in human coding sequence (0.074, 0.100 and 0.107; 26%). Closest-frequency recoding also left fewer codons far from native usage (20.3% shifted by more than 0.10, against 40.7% and 42.4%), although all three rules shared the same maximum shift of 0.433.

## Discussion

In this report we introduce an efficient tool for all-inclusive design of CRISPR/Cas9 modifications that combines multiple established tools with the latest technology (SMART) for efficient and precise genomic modification. Auto-SMART application provides comprehensive off-target evaluation delegated to CRISPOR and primer optimization delegated to Primer3, both linked directly from the result card. It provides a comprehensive suite of integrated applications from target selection to reagent ordering: edit selection, codon-usage-driven silent gap recoding, intron targeting under splice-signal protection, verified re-cut avoidance, and an HDR-allele preview that also seeds the genotyping primers. Precise design with the SMART approach requires concurrent consideration of the transcript frame, the cut geometry and the donor template which Auto-SMART achieves in a one stop application. We have performed extensive validation of annotations against an orthogonal protein database, an uncommon practice among CRISPR design tools, which generally accept whatever transcript they select. Notably, commercial HDR-donor design tools also incorporate blocking mutations, an established practice in repair-template design ^3,4^. However, none of these tools are in open access and also lack implementation of key design principles of Auto-SMART: verifying that each blocking change is amino-acid-silent before applying it, and declining to proceed when no silent change is available.

The current version of Auto-SMART is designed primarily for SMART editing with SpCas9 and NGG PAM sites. Although the platform identifies and ranks candidate guide RNAs, its on-target activity scores are computational predictions and should therefore be used to prioritize guides rather than as a substitute for experimental validation. Auto-SMART also relies on curated RefSeq and MANE transcript annotations when defining coding sequences and designing edits. Consequently, the transcript selected by the software may not always correspond to the isoform preferred for a particular biological application, and users should verify the selected transcript before proceeding.

Several editing strategies have not been incorporated into the current workflow. For example, Auto-SMART does not presently automate base-editing ^17^ or prime-editing ^18^ designs. Planned development will expand support to additional nucleases and PAM sequences, introduce dedicated base- and prime-editing workflows. Future versions are also expected to support batch processing and standardized export of design results, together with a more comprehensive tutorial containing representative inputs, outputs, and use cases.

In summary, by integrating guide selection, donor construction, silent recoding, and design evaluation into a single reproducible workflow, Auto-SMART substantially simplifies the implementation of SMART editing and CRISPR/Cas9 editing in general. This streamlined framework lowers the practical and technical barriers to using genomic editing for endogenous protein labeling and other precise modifications of genomic sequences.

## Methods

### Architecture

Auto-SMART is a static browser client served by a lightweight Python backend (Python 3.11). The client implements the interactive sequence viewer, the edit-operation state machine, SpCas9 guide scanning, SMART template assembly, re-cut prediction, the three-panel visualization, PDF export and Primer3 payload construction. The backend exposes a small API for gene and transcript lookup, genomic sequence retrieval, genome-to-taxonomy mapping, codon-usage proxying, on-target scoring; it serves static assets while blocking direct access to server-side files. The application is deployed on a managed cloud host.

### Input, target selection and transcript choice

Transcript models and genomic sequence are retrieved through the UCSC Genome Browser REST API ^9,19^. Gene and transcript lookups probe curated RefSeq tracks first (ncbiRefSeqSelect/MANE, then curated NM_/NR_) ^20,21^ before predicted models, ranking candidates by curated status and coding-sequence length so that the default transcript is the canonical, MANE-style isoform where available. Because SMART gap mutation is defined for protein-coding sequence, region selection is codon-aware: silent substitutions are applied only where the selection and gap can be read in frame. A selection lying entirely within one intron is routed to the intron mode described below, in which the gap is recoded without reference to a reading frame. To help users confirm they are editing the intended isoform, Auto-SMART compares the translated coding sequence of the selected transcript with the reviewed UniProt canonical protein ^22^ for the same gene and surfaces a warning when they differ, with one click to switch transcripts.

### Guide discovery, scoring and ranking

Auto-SMART scans a window around the target for SpCas9 5’-NGG-3’ PAMs on both strands and extracts 20-nt protospacers, annotating each with strand, sequence, PAM, predicted cut position and cut-to-edit gap length. On-target activity is estimated with the Doench 2016 Rule Set 2 model via the Azimuth implementation bundled with the server ^15^, scored from the standard 30-mer context (4 nt upstream + 20-nt protospacer + 3-nt PAM + 3 nt downstream). Candidates are ranked by a transparent predicted-efficiency heuristic that rewards predicted activity and penalizes longer cut-to-edit gaps. Writing T for the predicted on-target score (0-100) and G for the cut-to-edit gap in bp, the three design types share the base term 7 + 0.36 x T and differ only in how the gap is treated: SMART, 7 + 0.36 x T - 0.1 x G; SMART-RC/CT, 7 + 0.36 x T - 0.05 x G; and traditional, (7 + 0.36 x T) x exp(−0.0631 x G). The linear SMART form follows the published regression of SMART editing efficiency on predicted on-target score and cut-to-edit distance, and the C-terminal SMART-RC/CT design (used when an insertion sits immediately upstream of the stop codon) is penalized at half that rate, both because its gap is measured from the cut to the first base of the stop codon rather than to a user-chosen edit point and because of the higher knock-in efficiency reported for that geometry ^5^; a cut falling at or inside the stop codon is scored as a zero gap. A traditional template decays exponentially rather than linearly because it rebuilds the gap verbatim and so competes with the cut chromosome across the whole span; the decay constant corresponds to a half-life of ln(2)/0.0631, about 11 bp, so predicted efficiency falls to roughly a quarter of its gap-free value across one protospacer plus PAM. Each candidate links out to CRISPOR for genome-specific off-target analysis ^6,7^, with the currently selected assembly passed through automatically.

### SMART repair-template construction

For each top-ranked guide, Auto-SMART builds a repair template carrying the edit and, where required, synonymous substitutions across the gap between the edit and the predicted cut. The relevant gap is computed in genomic coordinates, converted to coding-strand display order, and overlapping codons are recoded by choosing, for each codon, the synonymous alternative whose frequency in the selected codon-usage table is closest to the original codon (minimizing perturbation of codon bias) while changing only bases inside the gap. This frequency-matching rule is deliberately distinct from codon optimization, which selects the most frequent synonymous codon ^23,24^. Homology arms are sized from the combined length L of the inserted/replaced sequence and the gap (40 bp per arm for L <= 120, L/2 for L <= 200, otherwise 100 bp), and the donor is assembled in the correct order of cut site, edit and coding direction. The server reports the raw donor sequence and whether it falls in a single-stranded (ssODN) or double-stranded length regime ^5^, and annotates the visualization with guide, PAM, cut site, gap, silent mutations, homology arms, the deletion scar / insertion / replacement block, and translated amino acids.

A traditional (non-SMART) repair template is offered as an explicit alternative for every editing application, through a second pair of controls in each design dialog. A traditional donor is a plain HDR template: the gap between the cut and the edit is copied verbatim from the reference, so the donor rewrites no base anywhere, and a residual on-target site, while still predicted and reported, cannot be blocked. Requesting a traditional template for an insertion immediately upstream of a stop codon overrides SMART-RC/CT, whose reconstruction across the cut-to-stop interval cannot by construction leave the gap verbatim, and the design falls back to the plain geometry. The choice is not offered for site-unspecific intron insertion, where a zero-length gap makes the two templates, and the two scoring formulas, identical.

### Automatic re-cut avoidance

After HDR, an edited allele can be re-cleaved by the same ribonucleoprotein if a cut-competent copy of the target survives ^3,4^. Auto-SMART models a surviving site as one that retains an intact NGG PAM and whose PAM-proximal seed (the 12 nt nearest the PAM) matches the guide within a strict mismatch threshold ^1,25^. When such a residual site is detected in the designed donor, the algorithm attempts to neutralize it by, in order of preference: disrupting the PAM with a synonymous codon substitution, a free base change at a position that is genuinely untranslated, or introducing at least two synonymous mismatches into the seed. Two details of the seed strategy matter for correctness. First, a candidate substitution is accepted only if it net-adds a mismatch against the guide; a naive “change N seed bases” heuristic can recode an already-mismatched base back toward the guide sequence and so increase re-cut risk while appearing to act. Second, because SpCas9 cuts bluntly 3 nt 5’ of the PAM ^1^, blocking substitutions are placed on the codons nearest that cut and only as many are introduced as are still needed, counting mismatches already present, so the donor is perturbed as little as possible and only where it matters for homology. Every candidate substitution is verified to be amino-acid-silent before it is applied: the server rebuilds the donor, re-reconstructs the edited allele, re-translates it and confirms that every changed codon still encodes the native residue. A base that the design synthesizes rather than copies from the genome, such as an inserted, replaced or point-mutated base, has no genomic coordinate and is therefore classified from the reading frame carried across the contiguous edited locus rather than from the transcript annotation. Where such a base is translated it is treated as coding, so that only a synonymous swap can reach it, and a stop codon placed by the design is treated as unrecodable rather than merely synonymous, because TAA and TAG both read as a stop. For a synthesized base the invariant verified is that blocking did not change what the donor already encoded, since a synthesized codon has no native amino acid to be compared with and a requested point mutation deliberately differs from one. Codons split across an intron, and any window it cannot verify, are rejected conservatively. Positions inside the protected splice zones of an intron, the three exonic bases flanking each splice junction, and the bases of a recognized insertion payload are removed from the candidate set before planning rather than being repaired afterwards. If no silent change can remove the risk, Auto-SMART reverts the attempt and displays an explicit “predicted re-cut” warning rather than silently altering the protein. The applied changes are highlighted in the donor and HDR panels so the user can audit them.

### Intron editing and splice-site protection

Introns are taken as the gaps between consecutive exons of the selected transcript, numbered biologically and reversed on the minus strand so intron and exon numbers agree. Terminal dinucleotides are classified in transcript orientation (GT..AG, GC..AG, AT..AC or non-canonical), which also verifies strand handling. Each intron is split into two protected zones and an editable interior: a 25 bp 5′ donor zone covering the U1 element, and a 60 bp 3′ acceptor zone covering the terminal AG, the polypyrimidine tract and a branch point 18–40 nt from the intron end. U12-type AT..AC introns receive a 30 bp donor zone, introns with unrecognized termini are protected at the wider default at both ends, and an intron too short to leave an interior cannot be edited.

Inside the interior, site-unspecific insertion enumerates only guides whose cut and full protospacer-plus-PAM footprint fall inside, sets the insertion point to the cut so the gap is zero, and ranks by predicted on-target activity; site-specific editing applies all four applications to a selected region, with the splice constraint acting on the union of cut site, edit footprint and gap. In both, the per-base veto extends three bases into each flanking exon and also governs gap recoding and the re-cut blocker.

Lacking a reading frame, the gap is recoded by complementing every base (A to T, C to G), which removes any internal protospacer or PAM while preserving GC content; frameshift checking is suppressed and a base picker replaces the residue row. Payloads are typed freely or drawn from a curated library (loxP, lox2272, lox5171, minimal 34 bp FRT, rox, piggyBac TTAA) that the re-cut blocker never alters. Inserts plus 30 bp of flanks are screened against donor, acceptor and branch-point consensus; hits overlapping changed bases are reported as advisory annotations.

### Codon usage and species support

Silent gap mutation and re-cut avoidance both require synonymous codon choice, which depends on the target organism’s codon usage. Auto-SMART uses bundled species-specific codon-usage tables where available and otherwise retrieves HIVE-CUTs/CoCoPUTs RefSeq codon-usage data for the assembly’s NCBI taxonomy identifier ^10,11^. If a species table cannot be retrieved, the server warns the user and falls back to a default table rather than silently changing design assumptions. The interface ships with common vertebrate and model-organism assemblies and also accepts custom UCSC assemblies when compatible transcript data are available.

### Output, export and privacy

Each design is presented as a result card: a summary header (guide number, sequence, PAM, strand, predicted on-target score, cut-to-edit gap, predicted SMART efficiency and codon-usage source), the raw donor sequence with a copy control, a Primer3 genotyping handoff, and three aligned interpretation panels (target locus, SMART donor template and locus after HDR). A “Change a gRNA” control cycles through lower-ranked candidates; a “Download SMART design” control exports a print-ready report; and a “Method description” link provides a concise methods paragraph. The Primer3 handoff passes a reconstructed HDR allele to Primer3 ^12–14^ with default constraints that keep primers away from the edit so that genotyping amplicons span the modification. Seeding Primer3 with the predicted edited allele rather than the reference matters in practice: a primer placed on the reference can fall inside the recoded gap or the insertion and so mismatch the very allele the assay is meant to detect. Usage logging records only operational metadata (timestamp, gene, transcript, assembly and editing application); submitted sequences and generated donor templates are never stored server-side.

### Benchmarking

Cross-species coverage and throughput were measured by running all four applications on 30 distinct genes in each of 20 assemblies (2,400 designs) against the deployed build, recording per-design wall-clock time. Transcript fidelity was assessed by translating the coding sequence of the transcript Auto-SMART selects and comparing it with the reviewed UniProt canonical protein ^22^ for the same gene, across 600 comparable genes per species. Re-cut behavior was quantified by sweeping coding windows across 36 human genes and classifying the top-ranked design in each window (1,152 designs). Guide-space availability was measured by enumerating every SpCas9 guide whose blunt cut falls within the coding sequence of 100 human genes (37,980 guides, all scored with Rule Set 2 ^15^) and asking, for 4,000 evenly spaced coding edit sites (40 per gene), how many guides and what maximum predicted activity are available at each allowed cut-to-edit distance. Codon-bias preservation was assessed by recoding each of the 59 human sense codons that have a synonymous alternative under three rules (Auto-SMART’s closest-frequency choice, codon optimization to the most frequent synonym, and random synonymous choice) and recording the absolute change in relative synonymous codon usage (|dRSCU|) ^26^, both unweighted across codon types and weighted by each codon’s occurrence in human coding sequence.

## Data availability

Example designs were produced from public genome and transcript resources queried by the web server (UCSC Genome Browser, UniProt, HIVE-CUTs/CoCoPUTs). All benchmark datasets underlying Figures 4, 6 and 7 are provided as Supplementary data (Supplementary Tables S1-S6).

## Supporting information

All benchmark datasets underlying Figures 4, 6 and 7 are provided as Supplementary data (Supplementary Tables S1-S6).

## Acknowledgements

We thank the developers of CRISPOR, Primer3, the UCSC Genome Browser, the HIVE-CUTs/CoCoPUTs codon-usage resources, UniProt and the Azimuth/Rule Set 2 guide-activity model, whose resources are used or linked by the Auto-SMART workflow.

## Author Contributions

C.Z. conceived and developed the Auto-SMART web platform, performed all in silico assays, analyzed the data, and wrote the initial draft of the manuscript. Y.C. tested the Auto-SMART web platform to identify improvements. K.A.M. supervised the study and revised the manuscript.

